# *Mycobacterium abscessus* persistence in the face of *Pseudomonas aeruginosa* antagonism

**DOI:** 10.1101/2024.09.23.614414

**Authors:** Rashmi Gupta, Martin Schuster, Kyle H. Rohde

## Abstract

Chronic bacterial infections are responsible for significant mortality and morbidity in cystic fibrosis (CF) patients. *Pseudomonas aeruginosa* (*Pa*), the dominant CF pathogen, and *Mycobacterium abscessus* (*Mab*) both cause persistent pulmonary infections that are difficult to treat. Co-infection by both bacterial pathogens leads to severe disease and poor clinical outcomes. To explore understudied interactions between these two CF pathogens, we employed culture- and molecular-based approaches. In a planktonic co-culture model, growth of *Pa* continued unimpeded, and it exerted a bacteriostatic effect on *Mab*. Strikingly, exposure of *Mab* grown in monoculture to cell-free spent supernatant of *Pa* resulted in a dramatic, dose-dependent bactericidal effect. Initial characterization indicated that this potent *Pa*-derived anti-*Mab* cidal activity was mediated by a heat-sensitive, protease-insensitive soluble factor of >3kDa size. Further analysis demonstrated that expression of this mycobactericidal factor requires LasR, a central regulator of *Pa* quorum sensing (QS). In contrast, ΔLasR *Pa* was still able to exert a bacteriostatic effect on *Mab* during co-culture, pointing to additional LasR-independent factors able to antagonize *Mab* growth. However, the ability of *Mab* to adapt during co-culture to counter the cidal effects of a LasR regulated factor suggested complex interspecies dynamics. Dual RNAseq analysis of *Mab-Pa* co-cultures revealed significant transcriptional remodeling of *Mab*, with differential expression of 68% of *Mab* genes compared to minimal transcriptional changes in *Pa*. Transcriptome analysis reflected slowed *Mab* growth and remodeling of carbon and energy metabolism akin to a non-replicating persister-like phase. A tailored *Mab* response to *Pa* was evident by enhanced transcript levels of genes predicted to interfere with alkylquinolone QS signals or provide protection against respiratory toxins and hydrogen cyanide. This is the first study to provide a transcriptome-level view of genetic responses governing the interplay between two important CF pathogens. This will provide insights into interspecies interaction mechanisms which can be targeted to disrupt their communities in a CF lung to improve patient clinical performance. Moreover, identification of a novel antimicrobial natural product with potent cidal activity against *Mab* will provide a chemical biology tool for identifying new drug targets in *Mab*.

## Introduction

Chronic microbial infections are responsible for morbidity and mortality in patients suffering with lung disorders, in particular cystic fibrosis (CF). CF is an autosomal recessive disorder characterized by a defect in cystic fibrosis transmembrane conduction regulator (CFTR) that regulates chloride movement in and out of the cell [1–3]. This results in improper salt balance leading to accumulation of thick, sticky mucus and defective mucociliary clearance from lung airways. This environment in which innate immune mechanisms are impaired is conducive for opportunistic pathogens to thrive, causing lung deterioration and poor quality of life of CF patients. Among the CF microbial community, notable bacterial pathogens include *Pseudomonas aeruginosa (Pa), Staphylococcus aureus, Burkholderia cenocepacia*, and nontuberculous mycobacteria (NTM) such as *Mycobacterium abscessus* (*Mab*) [4].

*Pa* is the dominant bacterial pathogen that can colonize lungs as early as within the first 5 years of life of a CF patient [5–9]. *Pa* produces several quorum sensing (QS) regulated virulence factors including proteases and secondary metabolites such as phenazines and hydrogen cyanide (HCN) detectable in CF sputum which facilitate colonization of CF airways [10–12]. These QS mediated factors help *Pa* gain dominance within the CF microbial community by restricting competition. The QS regulatory network of *Pa* is complex and comprised of four interconnected systems - *las*, *rhl*, *pqs* and *iqs* [13–16]. The first two systems are based on acylhomoserine lactone (AHL) signals, while the *pqs* system requires quinolone signals, and the fourth *iqs* responds to 2-(2-hydroxyphenyl)-thiazole-4-carbaldehyde signal molecules. The *las* system occupies the top position of the hierarchical QS system and is comprised of AHL signal synthase, LasI, and transcriptional regulator, LasR. This system positively regulates other systems of the QS circuitry; however these downstream pathways can also function independently of LasR under certain conditions [13, 15, 17–19]. LasR controls expression of numerous virulence genes and plays a crucial role in pathogenesis [13, 15]. Despite extensive study, the role of QS in CF infections remains perplexing. On one hand, *las* QS-regulated production of redox-active phenazine pigment pyocyanin causes pulmonary exacerbation and lung damage [20–22]. On the other hand, QS deficient LasR mutants are commonly isolated from CF patients and are responsible for enhanced neutrophil inflammation and disease progression [23–26].

*Mab,* a second bacterial inhabitant of CF airways, is a major threat due to an increased prevalence and a high level of inherent drug resistance [27–30]. Mutations in CFTR predispose CF patients to severe *Mab* infections [31]. *Mab* is a fast growing mycobacterium that colonizes and persists in the lungs of CF patients with a peak prevalence reported between age 10-15 [32]. *Mab* exhibits two different colony phenotypes - smooth and rough - and both are commonly isolated from CF patients [33–35]. Both phenotypes form biofilms, have similar antibiotic sensitivity and macrophage survival though the rough morphotype is considered more virulent [34, 36, 37]. Both *Mab* and *Pa* are versatile, multidrug resistant, CF pathogens which are difficult to treat.

*Pa* and *Mab* are isolated from CF sputum samples [28, 38, 39] and also share environmental (soil and water) [40] and host niches [41, 42]. Both *Pa* and *Mab* occupy the microaerobic respiratory zone in the lung [41, 42]. Sharing of lung spatial organization and horizontal gene transfer (HGT) from *Pa* to *Mab* [43] indicates co-existence of the two in lungs. Co-infection by these two opportunistic human pathogens is associated with pulmonary exacerbation, disease severity, and lung function decline [44–47]. Despite the obvious significant clinical relevance of the two bacterial pathogens, the nature of their interspecies interactions and potential impact on pathogenesis remains largely unexplored [1, 48–50]. Birmes *et al.* demonstrated quenching of quinolone QS signals of *Pa* by *Mab* when provided exogenously to *Mab* cultures [1]. Two other research groups examined the coexistence of the two in an *in vitro* dual-species biofilm model where *Pa* exerted a negative effect on *Mab* [48, 49]. However, *Pa-*mediated inhibition of *Mab* was reported to be contact-dependent and restricted to the solid surface biofilm model, with no inhibition in the planktonic co-cultures [48]. However, extensive analysis of *Pa* mutants lacking known virulence regulators (e.g. *las, rhl, pqs* QS systems), secretion systems, motility factors, and iron sequestration genes failed to identify the molecular mechanism responsible for this antagonism [48]. It was also noted that antibiotic-mediated targeted killing of *P*a favored *Mab* establishment in the dual-species biofilm [49]. It is imperative to interrogate interactions between these two human CF pathogens for a better understanding of their adaptive responses and impact on disease progression.

This study examined how *Mab* and *Pa* influenced each other through culture-based growth assays and molecular-based dual RNAseq analysis. This dual-species interaction study showed that *Mab* can persist when cultured alongside *Pa* but experiences substantial loss of viability when cultured in a cell-free *Pa* extract containing QS-dependent secreted factors. The non-cidal outcome in the co-culture and drastic changes at a molecular level in *Mab* strongly suggest that *Mab* employs adaptive mechanisms for survival and coexistence with *Pa*.

## Results and Discussion

### *Mab* antagonism in a planktonic co-culture with *Pa*

To gain an initial understanding of the direct interactions between *Pa* and *Mab,* we compared growth of individual species when present alone (monoculture) and in the presence of the other species (co-culture). A Δ*lasR* strain was also included to assess the role of LasR, a master transcriptional regulator of *Pa* QS in shaping interactions with *Mab*. As noted above, natural *lasR* mutants are frequently isolated from chronically infected CF patients, suggesting dysregulation of QS-regulated virulence factors may offer growth advantage to competitors [23]. Using the experimental setup illustrated in **Fig 1**, we first evaluated the growth of *Pa* and *Mab* in mono- and co-cultures by CFU enumeration. Growth of both WT and *ΔlasR Pa* strains remained unaffected in the presence of *Mab*. However, in contrast to the 2-3 log increase in CFU observed when *Mab* was grown alone, both WT and *ΔlasR Pa* exerted bacteriostatic antagonism of *Mab* (**Fig 2A, 2B**). These data indicate that the driver of bacteriostatic inhibition of *Mab* is a LasR-independent mechanism. This observation is in stark contrast to a recent report [48] where no *Mab* antagonism was observed in the planktonic co-culture with *Pa* [48]. The differences in media, strain and culture conditions likely contributed to this discrepancy. We utilized Luria-Bertani (LB) media as it is routinely used for *Pa* and *Mab* can also grow well in this growth media. The observed *Mab* growth arrest in the presence of *Pa* in this study indicates that *Mab* may prioritize survival over growth, potentially existing in a persister-like state in the co-culture.

**Fig 1.**
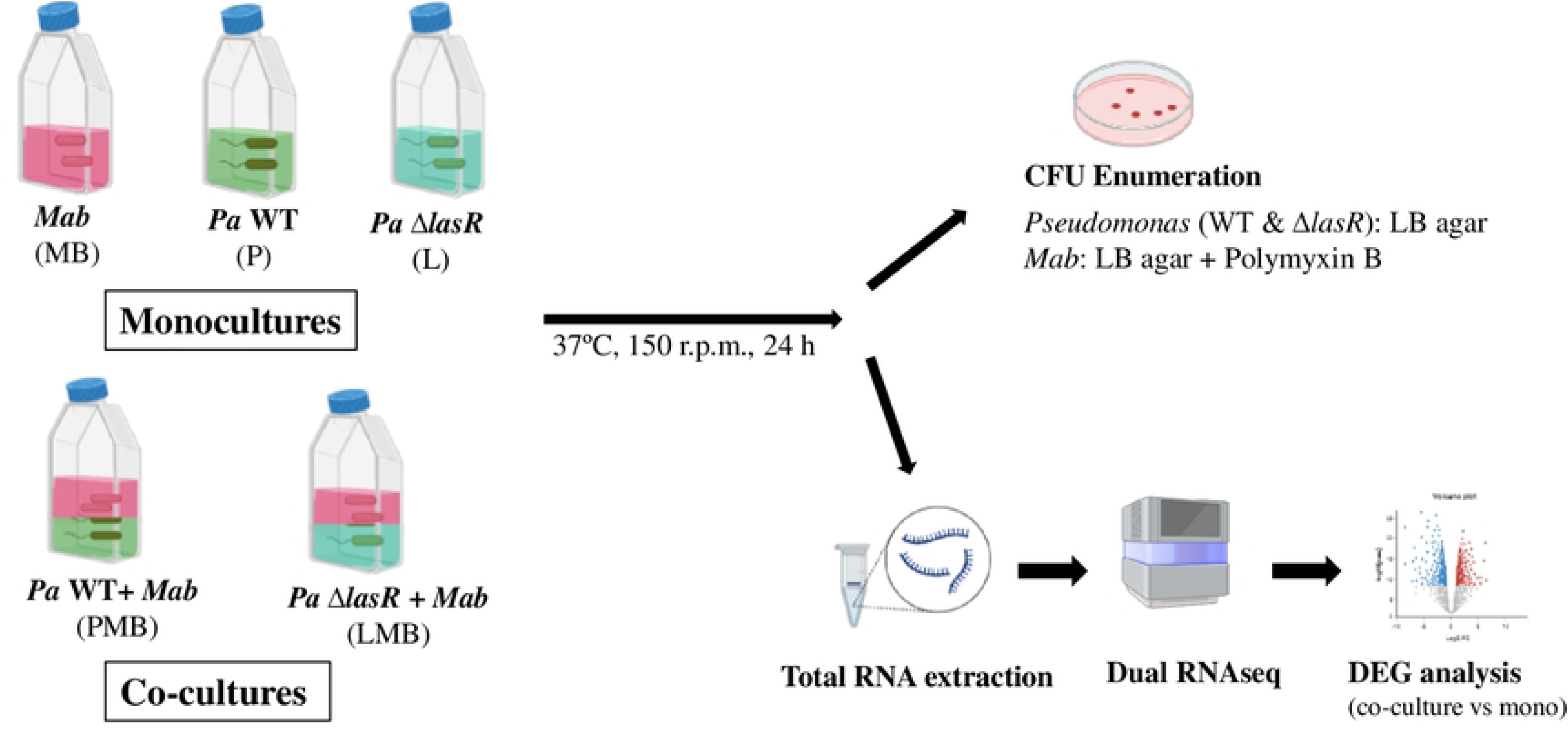
Planktonic co-culture model of *Mab-Pa* interactions. Monocultures of *Mab* (MB) and *Pa* strains (P and L) and co-cultures of *Mab+ Pa* WT (PMB) and *Mab*+ *Pa* Δ*lasR* (LMB) were grown for 24 h. *Pa* and *Mab* growth was evaluated by plating onto indicated selective media. The mono and co-culture samples were also processed for dual RNAseq analysis.

**Fig 2.**
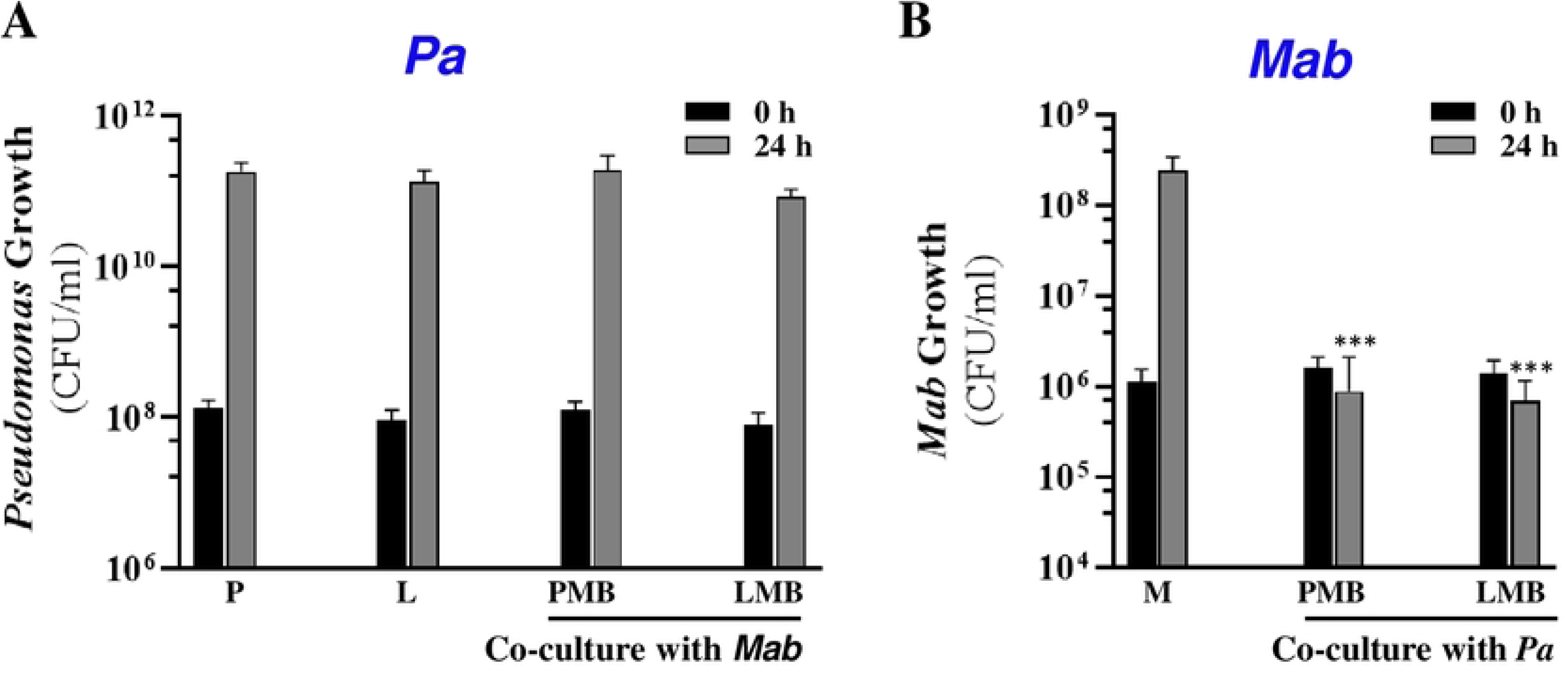
*Pa* antagonizes *Mab*. Growth (CFU/ml) of (A) *Pa* and (B) *Mab* strains when grown solo and in co-cultures at indicated times. P: *Pa* WT; L: *Pa* Δ*lasR*; MB: *Mab*; PMB: *Pa* WT+ *Mab;* LMB: *Pa* Δ*lasR* + *Mab.* *** denotes statistically significant difference (P<0.0001) from *Mab* monoculture after 24 h.

### Secreted *Pa* factor(s) are bactericidal toward *Mab*

Next, we investigated whether *Mab* inhibition by *Pa* in co-cultures was mediated by a contact-dependent mechanism or secreted factors. To address this, we examined the effect of *Pa* cell-free culture supernatant on *Mab* growth. When *Mab* was exposed to a range of concentrations (0%, 5%, 25% and 50% v/v) of *Pa* WT supernatant (sup), a dose- and time-dependent reduction in CFU was observed (**Fig 3A**). Exposure to 50% (v/v) *Pa* WT sup resulted in a 3-4 log decline in CFU compared to the control (no sup) within 24 h and complete sterilization within 48h. This indicates the presence of a highly potent mycobactericidal compound. However, this drastic cidal phenotype was lacking with the *Pa* Δ*lasR* supernatant, which caused only minor ∼1-log reduction in CFU at 48h (**Fig 3B**). Heterologous expression of the *lasR* allele from an arabinose inducible promoter in the *Pa* Δ*lasR* strain restored the cidal activity as indicated by the decline in *Mab* CFU and luminescence signal (**Fig 3C, S1 Fig**). This suggests that a LasR-dependent secreted factor (directly or indirectly regulated) mediated the potent cidal activity against *Mab.* We designated this cidal factor as SPAM (**S**ecreted ***P****seudomonas* **A**ntagonist of ***M****ab*) throughout the rest of this study.

**Fig 3.**
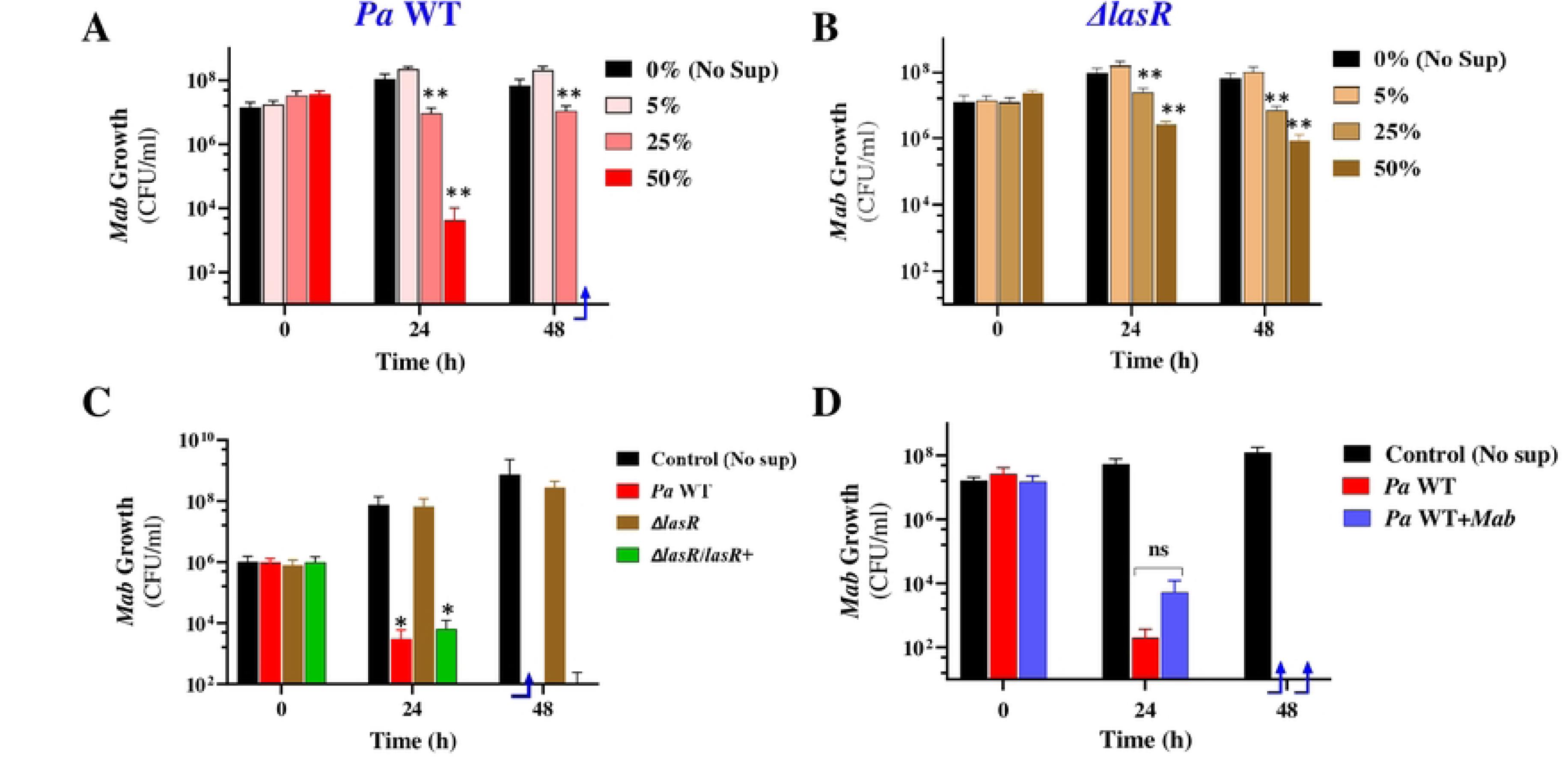
*Mab* viability in spent *Pa* supernatants. *Mab* growth (CFU/ml) in media supplemented with (A) *Pa* WT and (B) *Pa* Δ*lasR* culture supernatants at 0%, 5%, 25% and 50% concentration (v/v) (C) *Mab-lux* growth in 50%v/v spent *Pa* supernatants before after complementation of *lasR* mutation. A *lasR* allele was expressed from an arabinose inducible promoter carrying plasmid, pJN105. *Pa* WT and *Pa* Δ *lasR* strains carry the empty plasmid (D) *Mab* growth in sup obtained from *Pa* WT mono and co-cultures with *Mab*. Blue arrows indicate that no colonies were recovered. * P<0.01, ** P<0.001 from control, ns: insignificant (paired t-test).

The striking susceptibility of “naïve” *Mab* to killing by secreted *Pa* factors versus the tolerance of *Mab* when grown in co-culture with *Pa* suggest that interspecies interactions have either downregulated the production of cidal factors by *Pa* or rendered *Mab* more resistant to their effects, or both. To investigate these hypotheses, we explored the possibility of 1) downregulation of *Pa* secreted virulence factors and 2) *Mab* adaptation during co-culturing by evaluating *Mab* growth in 50% v/v spent supernatant from co-cultures as well. The fact that supernatants from *Pa* co-cultured with *Mab* exerted a cidal activity against *Mab* comparable to that seen with supernatant from *Pa* monoculture supports the conclusion that *Mab*-induced downregulation of SPAM is not occurring (**Fig 3D**). The remaining likely explanation is that *Mab* mounts adaptative responses to counter killing by *Pa* during their side-by-side growth. The molecular adaptation of *Mab* in response to *Pa* presence was explored through dual RNAseq analysis later in the study.

### SPAM Characterization

To characterize the potent SPAM factor, we assessed *Mab* growth in heat-denatured, protease-treated, and size-fractionated spent *Pa* supernatant. Heat inactivation of *Pa* (WT) supernatant largely abolished the cidal activity indicating the heat-labile nature of SPAM (**Fig 4A**). To determine whether SPAM is proteinaceous in nature, *Pa* sup was pretreated with proteinase K (Prot K) and viability of a *Mab-lux* reporter strain was evaluated through two readouts -luminescence and CFU. Prot K is a broad-spectrum serine protease that cleaves peptide bonds next to the carboxyl-terminal of aromatic and hydrophobic residues [51]. Two controls, untreated and Prot K treated media were included. The protease treatment did not eliminate the cidal activity of *Pa* supernatant (**Fig 4B, S2A Fig**). Both the untreated and Prot K digested cultures showed a ∼4-5 log decrease in *Mab* CFU after 48 h compared to controls (**Fig 4B**). These results strongly suggest that SPAM is not proteinaceous in nature, although we cannot definitively rule out a protein factor involvement because some *Pa* secreted proteins may resist degradation by Prot K digestion [51]. Size fractionation of supernatant indicated that the potent cidal SPAM factor is >3kDa in size as the retentate fraction killed *Mab* but not the flow-through fraction (<3kDa) (**Fig 4C, S2B Fig**). This characterization would eliminate small molecules like phenazines or respiratory toxins (e.g. HQNO) as prime candidates, unless they were encapsulated in outer membrane vesicles as reported by Vrla *et al.* [52]. Further investigation will be required to isolate and identify this LasR-controlled interspecies antagonist.

**Fig 4.**
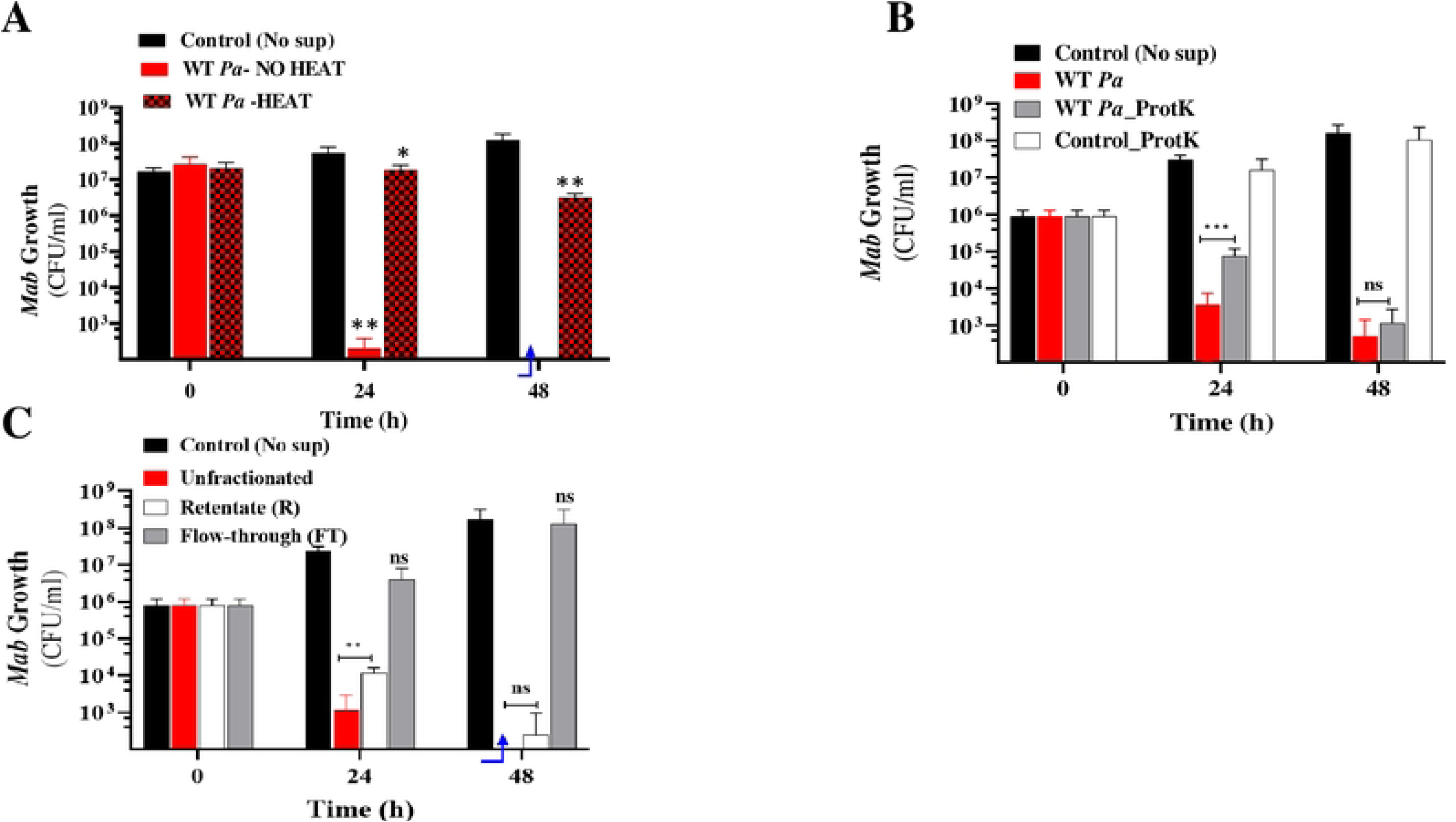
SPAM characterization. *Mab* growth (CFU/ml) in WT *Pa* supernatant (50% v/v) (A) with and without heat denaturation (B) proteinase K (ProtK) treated (C) Size fractionated (3kDa MWCO). The data is an average from at least three experiments. Note: For panel B and C, 10^1^ is the lowest dilution plated. Blue arrow denotes no detection of colonies. * P<0.01, ** p <0.001 from control, ***P=0.0001, ns: non-significant (two-tailed paired t-test).

Overall, our co-culturing and supernatant assays to investigate direct and indirect *Mab-Pa* interactions demonstrate a *Pa*-derived negative impact on *Mab* growth. Static growth of *Mab* when in physical contact with *Pa* appears to be mediated by either a *las* independent or a non-QS mechanism whereas cidal activity during indirect interaction with *Pa* is mediated via Las QS dependent secreted factors.

### *Mab* adaptation at a molecular level during co-culturing with *Pa*

To explore molecular cross talk between the two pathogens during co-culturing, we carried out dual RNAseq analysis and compared the changes in gene expression in both *Pa* and *Mab* in a co-culture vs a corresponding monoculture. We achieved a sequencing depth of 20-80 million total reads per sample meeting the minimum (10^6^) reads requirement for a comprehensive differentially expressed genes (DEGs) analysis [53, 54]. The differentially regulated *Mab* and *Pa* genes are listed in **S2 Table** and depicted with heat maps and Venn diagrams (**Fig 5**). Notably, the *Mab* transcriptome exhibited much more dramatic changes than *Pa* during co-culture of these two CF pathogens. More than 68% of the coding genes of *Mab* (3359/4920) were differentially regulated in the presence of WT *Pa*. Out of a total of 3359 *Mab* DEGs, 1800 were up-regulated and 1559 were down-regulated whereas a mere 275 and 226 *Pa* genes showed up and down-regulation, respectively (**Fig 5A, 5D**).

**Fig 5.**
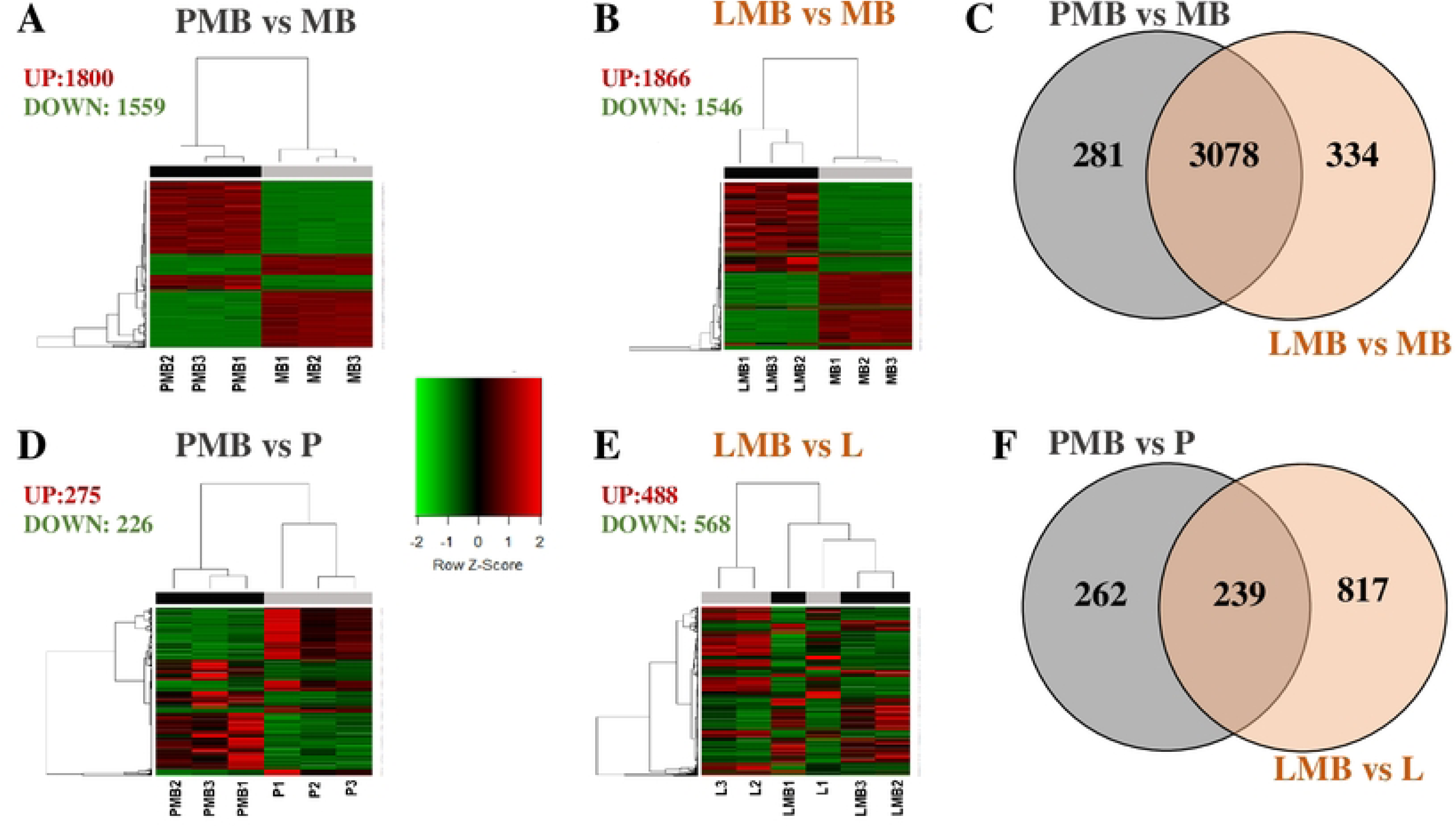
Transcriptomic responses of *Mab* (A, B and C) and *Pa* (D, E and F). Differentially expressed genes normalized counts per million of three replicates are represented by heat map (A, B, D, E) and overlap in expression by Venn diagrams (C, F). P: *Pa* WT, L: Δ*lasR*, MB: *Mab,* PMB: WT *Pa*-*Mab* co-culture, LMB: Δ *lasR* -*Mab* co-culture.

The transcriptional profile of *Mab* and *Pa* genes did not show any striking differences when the Δ*lasR Pa* strain was employed. *Mab* showed altered regulation of 3412 genes with up-regulation of 1866 and down-regulation of 1546 when co-cultured with Δ*lasR* (**Fig 5B**). Like WT *Pa,* Δ*lasR* showed relatively modest changes with 488 and 568 genes up- or down- regulated, respectively (**Fig 5E**). The Δ*lasR* strain showed downregulation of signature genes including direct LasR-dependent genes (*lasI, lasR, lasA, apr, rsaL*), phz2 operon, *hcnAB, qscR, rsaL,* as well as LasR controlled *rhl*-system (*rhlR, rhlAB,*) and *pqs* system genes (*pqsH*) as expected with loss of this QS signal receptor (**S2 Table**).

Overall, the extensive transcriptional reprogramming of *Mab* during co-culture with *Pa* reflects adaptive responses that likely contributed to *Mab’s* ability to withstand potentially cidal *Pa* secreted factors, whereas *Pa* appears to remain relatively undeterred during this rather one-sided interspecies interaction.

### Co-culture with *Pa* triggers slow-growing persister-like state in *Mab*

Persistence, which is characterized by slow or nonreplicating state, leads to antibiotic tolerance requiring prolonged treatment and high chances of treatment failure and recurring infections. Slow growth is marked by down-regulation of DNA replication, transcription and translation machinery, cell-division processes, electron transport, ATP-proton motive force and other cell processes to conserve energy [55]. Differential gene expression analysis between *Mab-Pa* co-cultures vs *Mab* monoculture revealed decreased carbon flux into metabolic pathways, reduced respiration, and energy metabolism (**S3 Table, S3 Fig**). Genes involved in cell division (*ftsZ*, *fts*K, *fts*X, *fts*E, *fts*H, and *ftsW*), DNA replication (*dnaA*, *dnaN*, *gyrA, gyrB*), and genes involved in transcription and translation (*fusA*, *tuf*, *tsf*, *rpoA, rpoB)* were repressed in the presence of *Pa.* A non-functional LasR did not change this response as *Mab* co-cultured with the Δ*lasR* exhibited a similar transcriptomic profile. There was also down-regulation of type-I NADH dehydrogenase encoded by *nuoA-N* genes, succinate dehydrogenase (*sdhABC*), cytochrome oxidases, and F_0_-F_1_ ATP synthase genes (*atpA-E*) (**Fig 6, S3 Table)**. This transcriptional response is akin to changes observed in *Mab* and in *M. tuberculosis* in its non-replicating persister phase [56, 57]. Entry of *Mab* into a non-replicating state could be a way to endure hostile conditions generated by *Pa* under co-culture conditions. This slow-growing state could have clinical implications, by contributing to the disconnect between *in vitro* and *in vivo* antibiotic susceptibility and to the ineffectiveness of current treatment regimens.

**Fig 6.**
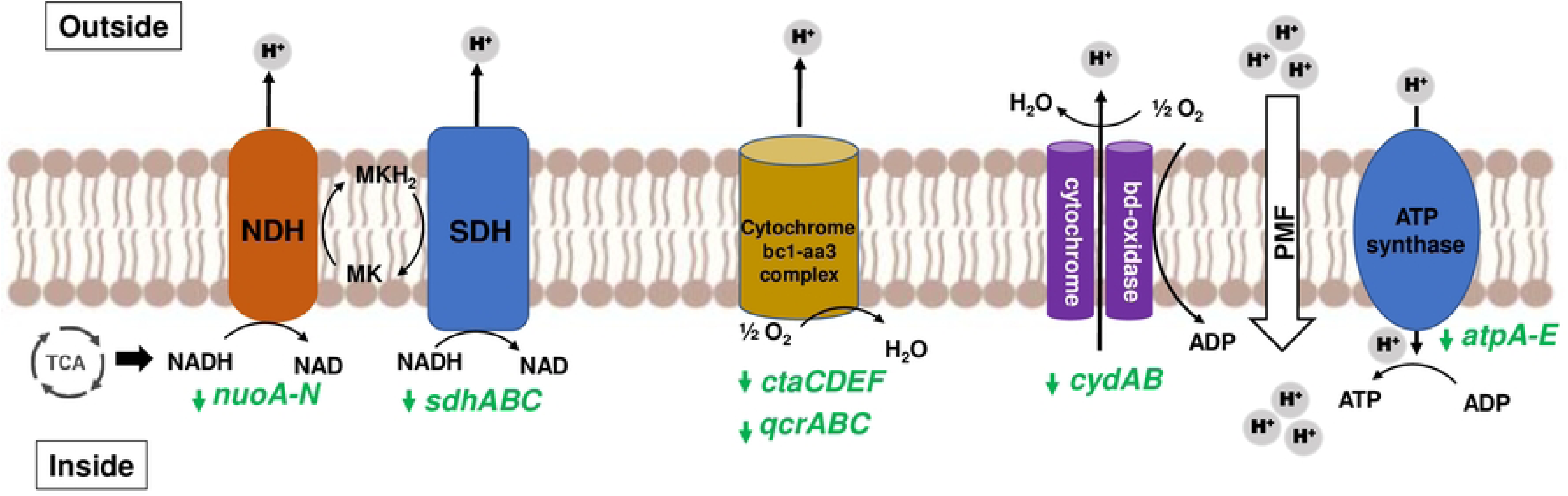
Slowing of *Mab* respiration during co-culturing. Mycobacterial NADH from TCA cycle enters mycobacterial electron transport chain (ETC) and supplies electrons to NADH dehydrogenase (NDH) and succinate dehydrogenase (SDH) to reduce menaquinone (MK) to menaquinol (MKH_2_). Cytochrome oxidases (bc1-aa3 complex and bd-type) oxidize back MK by transferring electrons to oxygen. Proton motive force (PMF) generated due to proton translocation is utilized by ATP synthase to produce ATP. Genes encoding ETC components are represented in green, and downward arrow denote downregulation.

### High induction of HGT elements in *Mab* during co-culture

The *Mab* genome contains a 23-Kb plasmid, pMAB23, and several other gene clusters predicted to be acquired by horizontal gene transfer (HGT) from closely related mycobacterial species and from non-mycobacterial community [43]. Strikingly, mobile elements, prophage and prophage-like elements and other HGT acquired genes including those putatively derived from *Pseudomonas* sp. were highly upregulated, with some having a log_2_ FC as high as 10. All 22 genes (MAB_p01 to MAB_p22c) encoded on the pMAB23 plasmid including the essential gene, *repA* (MAB_p16c), and putative arsenate resistance conferring genes were induced (**Fig 7, S4 Table**). In addition, several *Mab* genes in a 61 kb prophage region [58] (MAB_1723 to MAB_1833) exhibited strong transcriptional response in the presence of *Pa* (both WT and the *lasR* mutant). Three-prophage like elements [43] and 17-gene clusters acquired from the non-mycobacterial community were also differentially regulated. Out of the 17-clusters, 13 showed upregulation and only two, involved in degradation of phenylacetic acid (MAB_0899c-0911) and DNA (MAB_1093c-1098), respectively were down-regulated (**Fig 7, S4 Table**). The increased expression of mobile elements, prophage and other HGT genes in *Mab* under co*-* culture conditions suggest functional importance beyond being just passive elements in the genome. Increased expression of prophage genes is associated with survival under certain environmental conditions such as low pH in *M. avium* [59]. A transcriptomic response including higher expression of transposable elements, plasmid and viral-borne genes has been reported in other organisms such as slow-growing yeast cells for driving diversity for future stressful conditions [60].

**Fig 7.**
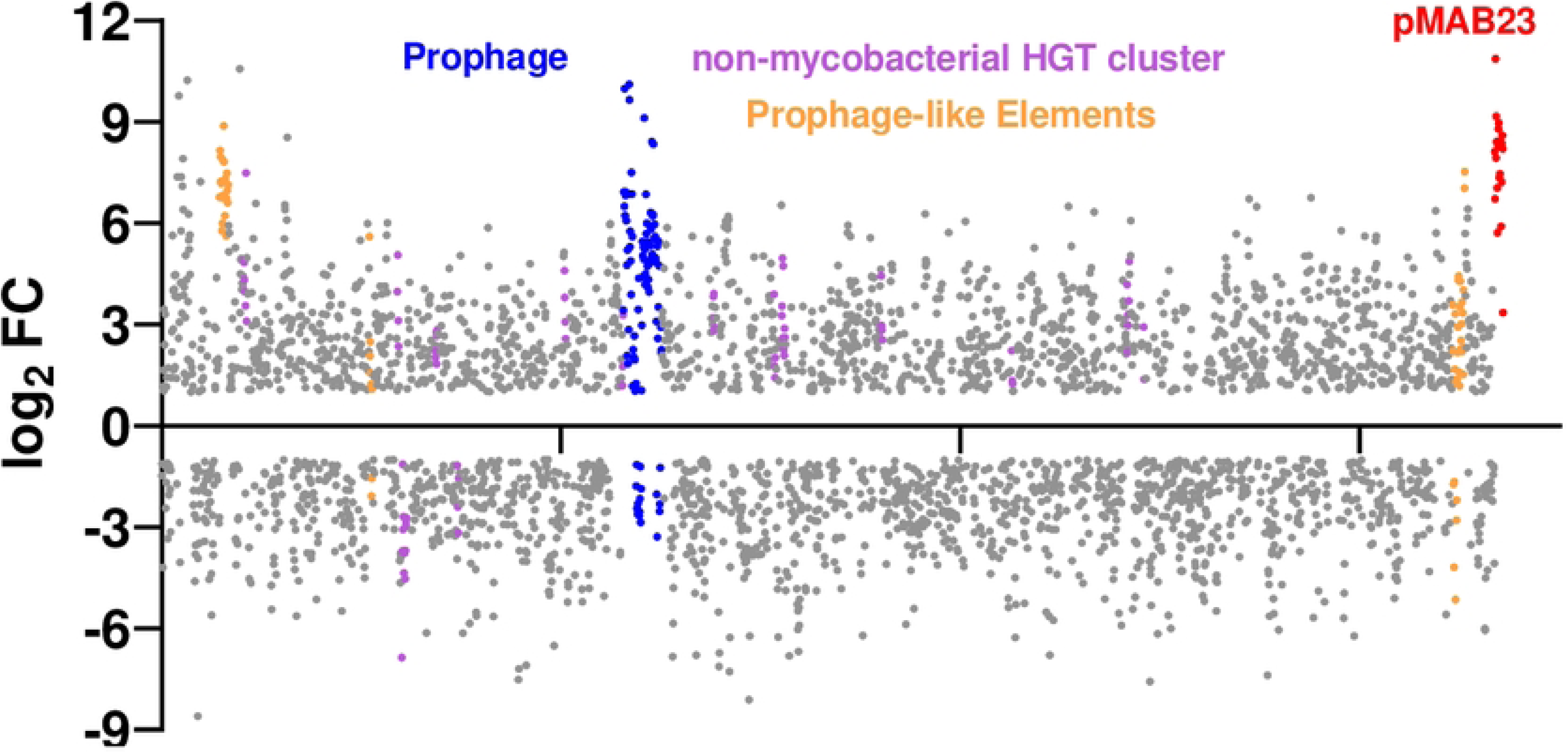
Up regulation of *Mab* HGT elements in a co-culture. A scatter plot showing differential gene regulation in *Mab* (PMB vs MB). Each dot represents a differentially regulated gene. The colored dots are the putative HGT acquired genes.

### Remodeling of central carbon metabolism of *Mab* in presence of *Pa*

One of the major transcriptional responses noted in *Mab* when co-cultured with *Pa* was in central metabolic pathways. RNAseq data seem to reflect a switch in carbon flow from sugars to fatty acids during co-culturing, with downregulation of glycolysis, gluconeogenesis, and Pentose Phosphate Pathway (PPP) pathways. The key enzymes involved in these pathways such as NADP-dependent dehydrogenase, aldolase, isomerases, and ATP-generating phosphoglycerate kinase, fructose 1,6-bisphoatase (*glpX*) and PEP carboxykinase (*pckA*) (**Fig 8**, **S5 Table**) were downregulated. This indicates reduced energy and biosynthetic precursors demand, aligning with the *Mab* phenotypic shift from replicating to a slow growing phase. Along these same lines, energy-consuming anabolic pathways were downregulated while lipid catabolism increased. There was an upregulation of lipases and fatty acid β-oxidation genes, along with decreased fatty acid synthesis (**S5 Table**), which suggest lipid breakdown for membrane biosynthesis or storage as triacylglycerol (TAG) for persistence and future energy source. The elevated transcripts of methylcitrate cycle (MCC) genes (*prpDCB*) and methylmalonate-semialdehyde dehydrogenase encoded by *mmsa* (log_2_ FC: 4) indicate propionyl CoA accumulation and subsequent detoxification [61, 62] (**Fig 8, S5 Table**). Reduced expression of propionyl-CoA carboxylase (*accD5*), methylmalonyl-CoA mutase (*mutB*) genes reflect preferential utilization of MCC over methyl malonyl (MM) for propionyl detoxification (**S5 Table**). Upregulation of mycolic acid synthesis genes (MAB_2028-MAB_2031 and beta-keto acyl synthases) hints at the incorporation of acetyl and propionyl CoA into mycolic acids. Key enzymes, *kasA* and *kasB* with a role in elongation of meromycolate chain showed upregulation (**S5 Table**). In addition, upregulation of triacylglycerol synthases (*tgs*) indicates TAG synthesis. Three out of 7 *tgs* genes encoded in the *Mab* genome (MAB_0854, MAB_1278, and MAB_2964), exhibited increased expression. This differential induction of *tgs* is also noted in *M. tuberculosis* under different inducing environments. Out of 15 *tgs* encoded in *Mtb*, *tgs1* was upregulated under hypoxia/nitric oxide stress condition whereas *tgs2* was induced by low pH [56, 63]. TAG accumulation is a hallmark of persisting *Mtb* which aids in drug tolerance and survival in the host [56].

**Fig 8.**
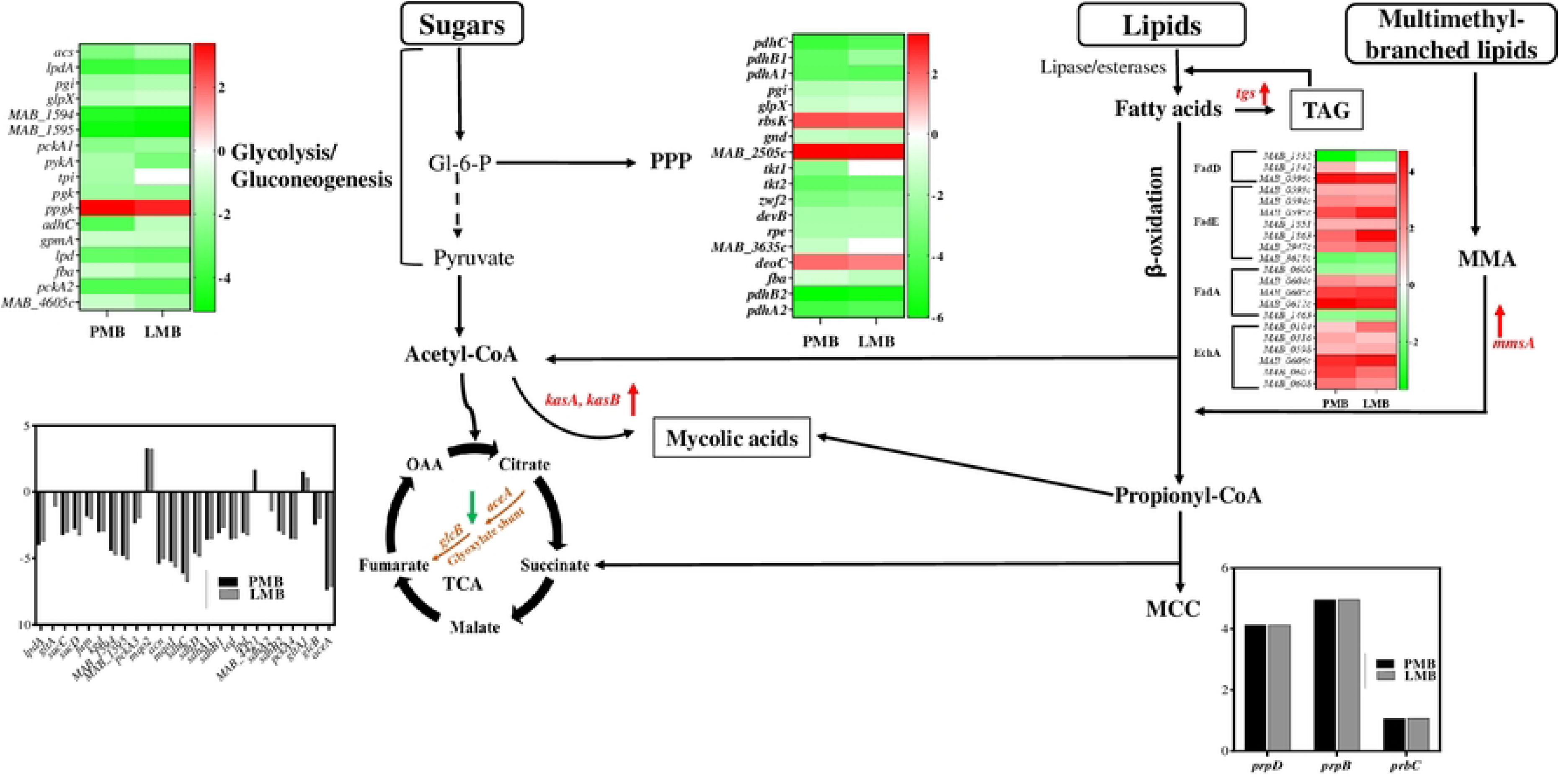
Carbon flow switch from sugars to fatty acids during co-culturing. The figure depicts an integrated view of central carbon metabolic pathways with differential expression of associated genes *via* heat maps, bar graphs, and by colored up and down arrows. Abbreviations: PPP; pentose phosphate pathway, TCA; tricarboxylic acid cycle, OAA; oxaloacetate, MCC; methyl citrate cycle, TAG; triacylglycerol, MMSA; Methylmalonate semialdehyde. Green and red arrow denotes down and upregulation, respectively. Carbon flow shunt to mycolic acids and TAG storage (boxed).

Based on the transcriptomic data, we speculate that excess fatty acids are channeled into intracellular lipids accumulation in the form of triacylglycerols (TAG) which can be hydrolyzed back by lipases when needed. TAG synthesis serves as a storage sink for persistence and survival under stress such as co-culture conditions. Resource limitation and competition during co-culture, perhaps enhanced by the lack of ample available sugars and other catabolizable carbon sources in LB medium [64], may be one driver of fatty acid catabolism and rerouting of carbon flow towards storage and alterations in cell wall composition to maintain cell viability and integrity. Similar carbon flow re-routing has been noted during *Mtb* infection [65, 66] and has implications in persistence and drug tolerance. The phenotypic growth arrest and persister-like metabolic remodeling may also reflect the impact of respiratory toxins and other antimicrobial metabolites secreted by *Pa*. *S*ecreted *Pa* factors like HCN, pyocyanin, and quinolone N-oxides inhibit *Staphylococcus aureus* growth by blocking respiration, leading to a small-colony persister phenotype [67].

### Co-culturing of *Mab* and *Pa* triggers competition for Fe

Culturing *Mab* and *Pa* together forces them to compete for common essential nutrients including Fe to replicate and survive. *Pa* utilizes several strategies to acquire iron under infection conditions including production and secretion of siderophores [68, 69]. Transcriptional profiles from dual RNAseq showed upregulation of siderophore-mediated iron-uptake, and downregulation of iron-storage in both bacterial pathogens, indicative of an Fe-starvation response (**Fig 9, S6 Table**). This pattern of altered expression of iron-responsive genes in both *Pa* and *Mab* in our co-culture model is indicative of competition between the interacting pathogens for Fe and likely other essential nutrients.

*Pa* produces two extracellular Fe chelating siderophores called pyoverdine and pyochelin with high and low affinity to Fe, respectively. Pyochelin biosynthesis genes, *pchA* and *pchD* and the gene cluster involved in synthesis and maturation of the major siderophore, pyoverdine (*pvdQAPMNOFEDJIHLG*) showed elevated expression in WT *Pa* when co-cultured with *Mab* (**Fig 9, S6 Table**). Besides synthesis, genes involved in secretion of pyoverdine into the extracellular environment via an ATP-dependent *pvdRT-ompQ* also had elevated expression. The transcript levels of the regulatory gene *pvdS*, encoding an iron-starvation sigma factor known for positive regulation of pyoverdine synthesis, was also enhanced. Downregulation of *bfrB* (bacterioferritin), which is involved in Fe storage, is also consistent with Fe-limiting conditions. Siderophore-mediated Fe uptake and storage genes were not differentially regulated in *Pa* in the Δ*lasR Pa-Mab* coculture (**Fig 9, S6 Table**) as expected because siderophore production is a QS regulated phenomenon [70–72].

**Fig 9.**
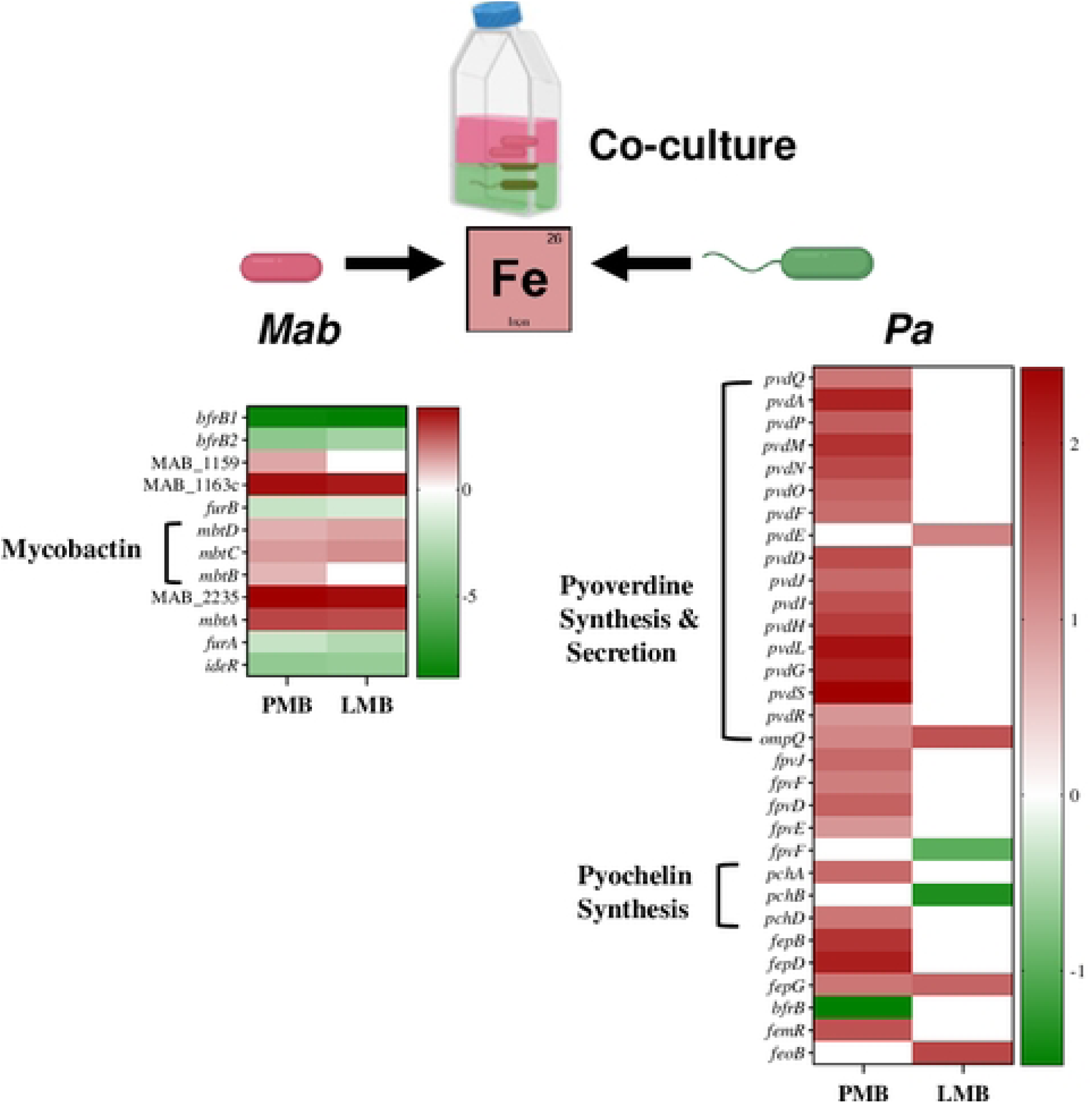
Competition for Fe among *Mab* and *Pa*. The heat map shows up and down regulated iron-responsive genes of *Mab* and *Pa* (WT and Δ*lasR*) during co-culture.

Similar to the Fe-limited transcriptional profile seen for *Pa* in co-culture, *Mab* also upregulated genes associated with iron uptake and downregulated iron storage genes. Genes encoding for the synthesis of the mycobacterial siderophore, mycobactin *(mbtDCBA)* showed higher expression whereas iron-dependent regulators (IdeR, FurA, and FurB) and the two putative Fe storage associated bacterioferritin B genes (MAB_0126, and MAB_0127) were downregulated (**Fig 9, S6 Table).**

The iron sequestering response from co-culture partners (*Pa* WT and *Mab*) indicates competition and survival efforts. Understanding Fe competition is vital for studying bacterial interactions, pathogenesis, and evolution of microbial communities. The adaption of various Fe –uptake strategies by both *Mab* and *Pa* in response to the co-culture environment is reminiscent of the human host infection conditions where the scarce availability of iron forces pathogens to adapt to survive and establish a successful infection. Effective consumption of limited nutrients by pathogens in CF lung will influence interaction dynamics and shape disease progression.

### Interspecies interaction between *Mab* and *Pa* alters virulence factor expression

Dual transcriptomic profiling revealed that the well-known virulence factors of *Pa* including both surface-associated and secreted factors did not show notable up or downregulation except for pyocyanin and siderophores (**S7 Table**). Our *in vitro* assays with *Pa* spent supernatant from mono and co-cultures also indicated no significant *Mab*-mediated downregulation of *Pa* secreted virulence factors.

However, in *Mab*, the presence of *Pa* triggered increased expression of several virulence factors such as MmpL/MmpS family lipid transporters, lipoproteins, and drug resistance associated genes involved in efflux and target modification (**S7 Table**). Several monooxygenases/dioxygenases implicated in ring cleavage of aromatic compounds were upregulated indicating *Mab* potential to inactivate *Pa* aromatic compounds such as pyocyanin and QS signals (**S8 Table**). Seventeen out of twenty-three dioxygenases present in *Mab* genome were up-regulated including the *aqd* cluster, MAB_0301-MAB_0303, known to degrade *Pa* AQ signals [1, 73] (**S8 Table**). Also, elevated expression of cyanate hydratases encoded by MAB_0054c and MAB_2545c hints at *Mab* upping its defense against HCN produced by *Pa* [74, 75]. This tailored response of *Mab* to *Pa* virulence factors indicates *Mab* adaptation efforts to survive to coexist with *Pa* in similar niches within CF lung.

## Conclusion

In this study, we exploited both culture–based and dual RNA transcriptomics strategies to explore interactions between *Pa* and *Mab*, two difficult-to-treat pathogens that commonly afflict CF patients. A handful of prior studies utilizing rudimentary biofilm models [48, 49] have reported *Pa-*mediated antagonism of *Mab,* although the molecular mechanisms underlying this interaction remain unknown. In contrast to the recent findings of Idosa *et al.*[48], this study is the first observation of *Pa* antagonism of *Mab* in a planktonic co-culture model, with *Pa* exerting a strong bacteriostatic effect on *Mab*. We also discovered the presence of soluble secreted factor(s) in *Pa* supernatants, which we dubbed SPAM, that exhibited potent bactericidal activity against *Mab.* Initial efforts to characterize SPAM indicate a protease-resistant, heat-labile factor of >3Kda whose expression is dependent on the LasR QS system. Isolation and identification of this natural product antimycobacterial agent, which is beyond the scope of the current study, may yield a valuable starting point for development of new antimicrobials and a chemical biology tool for identification of novel drug targets in *Mab.* The ability of *Mab* to withstand SPAM-mediated killing and persist alongside *Pa* in the co-culture model serves as evidence that interspecies interactions induce alternations in *Mab* that facilitate persistence. To gain genome-wide insight into global transcriptional changes triggered by *Pa-Mab* interactions, we conducted what is to our knowledge the first dual RNAseq analysis of these CF pathogens. *Mab* was notably more strongly impacted than *Pa* (based on number of differentially expressed genes), in line with *Pa’s* dominant role as a “bacterial bully” in its interactions with other species. In addition to *Mab* genetic signatures consistent with persistence and growth arrest we observed, there was evidence of adaptative responses to thwart known antimicrobial effectors of *Pa*. This valuable dataset provides novel insights into *Pa-Mab* interspecies interactions, however, much more work is needed to understand how these interspecies dynamics impact clinical outcomes and treatment. The apparent ability of *Mab* to withstand potentially bactericidal effectors secreted by *Pa* is likely critical for *Mab* to co-exist with *Pa* during chronic infections of vulnerable patients with CF or other pulmonary conditions like COPD. The well-documented evolution of *Pa* during chronic infection to yield variants lacking functional LasR QS could dramatically alter the dynamic interplay with other species like *Mab*. Our discovery of a LasR-dependent bactericidal antagonist of *Mab* would suggest this loss of LasR signaling in *Pa* at late stages of infections may give *Mab* the upper hand. Finally, the high-level drug resistance and distinct drug susceptibility profiles of *Pa* and *Mab* already makes treatment of polymicrobial infections challenging. The *Pa-*induced persister-like phenotype of *Mab* reflected in transcriptional signatures raise the possibility that interspecies interactions may impact antibiotic tolerance and treatment efficacy.

## Materials and Methods

### Bacterial strains and Reagents

The bacterial strains *Pa* WT (PAO1 Ochsner) [76], Δ*lasR* (in PAO1 Ochsner background) [77], and *Mab* 390S [36, 78] and *Mab-lux* [79] were used in this study. For routine culturing and maintenance, Luria Bertani (LB) was used for both *Pa* and *Mab* strains. For selection of *Mab* for species-specific CFU enumeration from co-cultures, LB agar supplemented with 200U/ml of antibiotic polymyxin B sulfate salt (Millipore sigma) was used. No selective media was necessary for *Pa* as it forms colonies after 24 h unlike *Mab,* which takes 3-5 days.

### In vitro co-culturing

A planktonic co-culture assay was used to investigate direct interactions between *Pa* and *Mab*. LB-Tween 80 (0.05%) media was inoculated from a freshly streaked single colony of *Pa* strains (WT and the *lasR* mutant) and *Mab* culture was started from a freezer stock. The bacterial cultures were appropriately diluted to set-up mono and co-cultures in 10 ml LB-Tween in T25 flasks. The starting OD of *Pa* and *Mab* was set at 0.05 and 0.15, respectively as used previously to account for growth rate differences [1, 80]. Three monocultures: WT (P), Δ*lasR* (L), and *Mab* (MB) and the two co-cultures -*Mab*+WT *Pa* (PMB), *Mab*+Δ*lasR* (LMB) were incubated at 37°C with shaking at 150 r.p.m. for 24 h. After 24 h, 10-fold serial dilutions were prepared in PBS media and spot-plated (5 µl) onto LB agar and LB agar supplemented with polymyxin B (200U/ml) for growth of *Pa* and *Mab,* respectively for colony enumeration.

### *Mab* growth in *Pa* supernatant

*Mab* growth was assessed in spent supernatants of WT (P) and the isogenic Δ*lasR* (L) strains of *Pa* to examine the effect of *Pa* secreted effectors. The *Pa* strains were grown overnight in LB-Tween media at 37°C, 150 r.p.m. shaking from a freshly streaked isolated colony. The cultures were diluted to OD_600_ of 0.05 and grown for 24 h. Thereafter, cultures were centrifuged (4000 r.p.m., 30 min., 10°C) and filtered through 0.22 µm pore size cellulose acetate filters (Corning, 43020). The filter-sterilized supernatant was plated (100 µl) onto LB agar plates to ensure that no *Pa* cells were present in the filtrate. *Mab* cultures were set at an OD_600_ of 0.15 in 10 ml LB-Tween (T25 flasks) with 0%, 5%, 25%, and 50% v/v, of *Pa* spent monoculture supernatant. The cultures were incubated at 37°C with shaking (150 r.p.m.) and after 0, 24, and 48 h 10-fold serial dilutions were spot-plated (5µl) onto LB agar for each time point. The CFU/ml data is an average from three biological experiments each with 4 technical replicates. The same assay was performed with 50% v/v supernatants derived from *Pa-Mab* co-cultures as well.

### *lasR* complementation

To achieve complementation, low-copy-number plasmids, pJN105 [81] and pJN105.*lasR* (pJN105.L) [82] were transformed into chemically competent *lasR* mutant cells as described [83]. Plasmid pJN105 served as an empty vector control whereas pJN105.*lasR* expressed *lasR* from an arabinose -inducible ara-BAD promoter. The spent supernatant from *Pa* strains (WT/pJN105, Δ*lasR/*pJN105, and complement Δ*lasR*/*lasR^+^*) was harvested as described earlier except the cultures were induced with 50mM arabinose during growth. *Mab* growth was evaluated in spent supernatants using a luminescence-based 96-well reporter assay [79]. Briefly, *Mab-lux* was grown in 50% v/v spent supernatant in a total well volume of 100 µl at a final OD_600_ of 0.01. The plates were incubated at 37°C and luminescence signal was measured after every 24 h for 2 days. An aliquot was taken for serial dilution and plating to determine CFU for the indicated times.

### Heat, Protease sensitivity and Size fractionation determination of SPAM

To gain insights into the heat stability, size and chemical nature of *Pa*-derived anti-mycobacterial factor known as SPAM, *Mab* growth was examined in heat-denatured, protease treated and size fractionated *Pa* supernatants. For heat treatment, filter-sterilized supernatants were heated at 95°C for 15 min. in a water bath. For protease treatment, cell-free *Pa* supernatant was incubated with 200 µg/ml proteinase K for 2 h at 37°C. Size fractionation of the spent supernatant was carried out by centrifuging through 3kDa MWCO filters (Millipore, SIGMA, UFC900308) for 20 min at 4000 r.p.m. Both retentate and flow-through fractionated samples were filter-sterilized using cellulose acetate, low protein binding syringe filters. *Mab* growth assessment in the heat-denatured supernatant was carried out in flasks as described in the supernatant assay while a high throughput microtiter plate assay described before in *lasR* complementation was adopted for proteinase K (Prot K) treated, and 3kDa size fractioned *Pa* supernatants. Controls including no supernatant and unfractionated/untreated supernatant were included in the assay. The data is an average from 3 biological experiments each with a total of 9 technical replicates.

### Total RNA extraction

Total RNA was isolated from mono and co-cultures involving *Pa* and *Mab* after 24 h of incubation at 37°C by using previously reported mycobacterial trizol-based RNA isolation technique [84] with modifications. *Mab* and *P*a cultures were mixed with RNA Protect reagent (Qiagen), mixed by inversion, and centrifuged (4300 r.p.m.) for 20 min. The supernatant was discarded, and pellet was resuspended in 1 ml RNA Protect reagent before storage at -80°C. The pellets were disrupted using a Mini-Bead Beater 16 (Biospec) 2x at max speed for (for 1 min followed by incubation on ice for 1 min. After this, steps were carried out at 4°C centrifuge as described previously using RNeasy kit (Qiagen) [84]. Integrity of isolated RNA was determined using Agilent RNA ScreenTape system.

### Dual RNAseq

Transcriptome analysis was carried out with three biological replicates of each sample (mono and co-cultures). Depletion of Ribosomal RNA, mRNA library preparation and sequencing were performed by SeqCenter (earlier MiGS). Library preparation was performed using Illumina’s Stranded Total RNA Prep Ligation with Ribo-Zero Plus kit and 10 bp IDT for Illumina indices. Around Total RNA (275ng) was ribodepleted using oligonucleotide sequences specific for *Pa* and *Mab*. These custom probes are listed in **S1 Table**. The samples were DNAase treated with Invitrogen DNAase (RNAase-free) and sequencing was performed (NextSeq2000, 2×50bp reads). Sequencing quality control and adapter trimming was performed with bcl-convert (v3.9.3) (https://support-docs.illumina.com/SW/BCL_Convert/Content/SW/FrontPages/BCL_Convert.htm). Read mapping was performed with HISAT2 [85] and read quantification was done using Subread’s featureCounts functionality [86]. Reads for *Pseudomonas* monoculture samples (P1, P2, P3, L1, L2, L3) were mapped to the *P. aeruginosa* PAO1 RefSeq genome NC_002516.2, *Mab* solo culture samples (MB1, MB2, and MB3) to the RefSeq genome GCF_000069185.1 (ASM6918v1), and co-culture samples (PMB1, PMB2, PMB3, LMB1, LMB2, and LMB3) were simultaneously mapped. Differential expression analysis was performed using edgeR’s Quasi-Linear F-Test (qlfTest) functionality against treatment groups. Differentially expressed genes (DEGs) were identified with a cut-off of log_2_ fold-change (FC) (-1 ≥1 to ≤1) and p-value < 0.05 (**S2 Table**). The KEGG pathway analysis was performed using limma’s [87] “kegga” functionality with genes with FDR<0.05 (adjusted p-value).

## Authors contributions

RG conceived the study, designed, and conducted experiments. RG carried out result analysis and drafted the manuscript. KR provided resources, technical inputs and helped in manuscript preparation. MS provided key reagents, suggestions and critical review of the manuscript.

## Data Availability

Sequencing data generated in this study have been deposited in the NCBI’s SRA database and can be accessed under BioProject ID PRJNA1154599. All other relevant data are within the manuscript and its <u>Supporting Information</u> files.

## Supporting Information

**S1 Fig.** *lasR* complementation. *Mab* growth measured by luminescence signal after complementation of *lasR* mutation. The assay was carried out in 96-well microtiter plates with 50% v/v spent supernatants of *Pa* strains. The data is an average from 3 experiments each with a total of at least 9 technical replicates.

**S2 Fig.** *Mab-lux* growth in *Pa* WT supernatants (50% v/v) after A) proteinase K (Prot K) treatment B) Size fractionation (3kDa MWCO).

**S3 Fig.** Functional classification of *Mab* DEGs in a co-culture with WT *Pa* vs monoculture. A bar chart showing select affected functional classes with up and down regulated (log_2_ ≥ 1 and FDR ≤ 0.05 genes.

**S1 Table.** *Pa* and *Mab* probes used for rRNA depletion

**S2 Table.** Differentially regulated genes in a co-culture versus monocultures (log_2_FC -1 ≥1 to ≤1, p<0.05).

**S3 Table.** Differentially regulated *Mab* genes related to slow growth when partnered with *Pa*

**S4 Table.** Differentially regulated, horizontally acquired *Mab* genes in a co-culture with *Pa*

**S5 Table.** Differentially expressed carbon metabolism genes in *Mab*

**S6 Table.** Differentially expressed iron-responsive *Mab* genes in a co-culture with *Pa*

**S7 Table.** Virulence related *Mab* DEGs in a co-culture with *Pa*

**S8 Table.** Differentially expressed mono/dioxygenases *Mab* genes in a co-culture with *Pa*

## References

1. Birmes FS, Wolf T, Kohl TA, Ruger K, Bange F, Kalinowski J, Fetzner S. Mycobacterium abscessus subsp abscessus Is Capable of Degrading Pseudomonas aeruginosa Quinolone Signals. Front Microbiol. 2017;8. PubMed PMID: WOS:000394980900001.

2. Elborn JS. Cystic fibrosis. Lancet. 2016;388(10059):2519–31. PubMed PMID: WOS:000388166300030.

3. Mall MA, Hartl D. CFTR: cystic fibrosis and beyond. Eur Respir J. 2014;44(4):1042–54. PubMed PMID: WOS:000342543700027.

4. Valenza G, Tappe D, Turnwald D, Frosch M, Konig C, Hebestreit H, Abele-Horn M. Prevalence and antimicrobial susceptibility of microorganisms isolated from sputa of patients with cystic fibrosis. J Cyst Fibros. 2008;7(2):123–7. Epub 20070813. doi: 10.1016/j.jcf.2007.06.006. PubMed PMID: 17693140.

5. Aebi C, Bracher R, Liechtigallati S, Tschappeler H, Rudeberg A, Kraemer R. The Age at Onset of Chronic Pseudomonas-Aeruginosa Colonization in Cystic-Fibrosis - Prognostic-Significance. Eur J Pediatr. 1995;154(9):S69–S73. doi: Doi 10.1007/Bf02191510. PubMed PMID: WOS:A1995RU72200019.

6. Banjar H, Ghawi A, AlMogarri I, Alhaider S, Alomran H, Hejazi A, et al. First report on the prevalence of bacteria in cystic fibrosis patients (CF) in a tertiary care center in Saudi Arabia. Int J Pediatr Adolesc Med. 2022;9(2):108–12. Epub 20210707. doi: 10.1016/j.ijpam.2021.07.001. PubMed PMID: 35663786; PubMed Central PMCID: PMCPMC9152558.

7. Bhagirath AY, Li YQ, Somayajula D, Dadashi M, Badr S, Duan KM. Cystic fibrosis lung environment and Pseudomonas aeruginosa infection. Bmc Pulm Med. 2016;16. PubMed PMID: WOS:000390208600001.

8. Pittman JE, Calloway EH, Kiser M, Yeatts J, Davis SD, Drumm ML, et al. Age of Pseudomonas aeruginosa Acquisition and Subsequent Severity of Cystic Fibrosis Lung Disease. Pediatr Pulm. 2011;46(5):497–504. doi: 10.1002/ppul.21397. PubMed PMID: WOS:000289510500012.

9. Van Daele SG, Franckx H, Verhelst R, Schelstraete P, Haerynck F, Van Simaey L, et al. Epidemiology of Pseudomonas aeruginosa in a cystic fibrosis rehabilitation centre. Eur Respir J. 2005;25(3):474–81. doi: 10.1183/09031936.05.00050304. PubMed PMID: 15738291.

10. Chambers D, Scott F, Bangur R, Davies R, Lim A, Walters S, et al. Factors associated with infection by Pseudomonas aeruginosa in adult cystic fibrosis. Eur Respir J. 2005;26(4):651–6. doi: 10.1183/09031936.05.00126704. PubMed PMID: 16204596.

11. Singh PK SA, Parsek MR, Moninger TO, Welsh MJ, Greenberg EP. Quorum-sensing signals indicate that cystic fibrosis lungs are infected with bacterial biofilms. Nature. 2000 Oct 12;407(6805):762-4. PMID: 11048725. Nature. 2000;(Oct 12;407(6805)):762-4. doi: doi: 10.1038/35037627. . PubMed Central PMCID: PMC11048725.

12. Jan AT. Outer Membrane Vesicles (OMVs) of Gram-negative Bacteria: A Perspective Update. Front Microbiol. 2017;8:1053. Epub 20170609. doi: 10.3389/fmicb.2017.01053. PubMed PMID: 28649237; PubMed Central PMCID: PMCPMC5465292.

13. Latifi A, Foglino M, Tanaka K, Williams P, Lazdunski A. A hierarchical quorum-sensing cascade in Pseudomonas aeruginosa links the transcriptional activators LasR and RhIR (VsmR) to expression of the stationary-phase sigma factor RpoS. Mol Microbiol. 1996;21(6):1137–46. PubMed PMID: WOS:A1996VJ51800003.

14. Lee J, Wu J, Deng Y, Wang J, Wang C, Wang J, et al. A cell-cell communication signal integrates quorum sensing and stress response. Nat Chem Biol. 2013;9(5):339–43. Epub 20130331. doi: 10.1038/nchembio.1225. PubMed PMID: 23542643.

15. Pesci EC, Pearson JP, Seed PC, Iglewski BH. Regulation of las and rhl quorum sensing in Pseudomonas aeruginosa. J Bacteriol. 1997;179(10):3127–32. doi: 10.1128/jb.179.10.3127-3132.1997. PubMed PMID: 9150205; PubMed Central PMCID: PMCPMC179088.

16. Wade DS, Calfee MW, Rocha ER, Ling EA, Engstrom E, Coleman JP, Pesci EC. Regulation of Pseudomonas quinolone signal synthesis in Pseudomonas aeruginosa. J Bacteriol. 2005;187(13):4372–80. doi: 10.1128/JB.187.13.4372-4380.2005. PubMed PMID: 15968046; PubMed Central PMCID: PMCPMC1151766.

17. Dekimpe V, Deziel E. Revisiting the quorum-sensing hierarchy in Pseudomonas aeruginosa: the transcriptional regulator RhlR regulates LasR-specific factors. Microbiology (Reading). 2009;155(Pt 3):712–23. doi: 10.1099/mic.0.022764-0. PubMed PMID: 19246742.

18. Diggle SP, Winzer K, Chhabra SR, Worrall KE, Camara M, Williams P. The Pseudomonas aeruginosa quinolone signal molecule overcomes the cell density-dependency of the quorum sensing hierarchy, regulates rhl-dependent genes at the onset of stationary phase and can be produced in the absence of LasR. Mol Microbiol. 2003;50(1):29–43. doi: 10.1046/j.1365-2958.2003.03672.x. PubMed PMID: 14507361.

19. Medina G, Juarez K, Diaz R, Soberon-Chavez G. Transcriptional regulation of Pseudomonas aeruginosa rhlR, encoding a quorum-sensing regulatory protein. Microbiology (Reading). 2003;149(Pt 11):3073–81. doi: 10.1099/mic.0.26282-0. PubMed PMID: 14600219.

20. Caldwell CC, Chen Y, Goetzmann HS, Hao Y, Borchers MT, Hassett DJ, et al. Pseudomonas aeruginosa exotoxin pyocyanin causes cystic fibrosis airway pathogenesis. Am J Pathol. 2009;175(6):2473–88. Epub 20091105. doi: 10.2353/ajpath.2009.090166. PubMed PMID: 19893030; PubMed Central PMCID: PMCPMC2789600.

21. Hunter RC, Klepac-Ceraj V, Lorenzi MM, Grotzinger H, Martin TR, Newman DK. Phenazine content in the cystic fibrosis respiratory tract negatively correlates with lung function and microbial complexity. Am J Respir Cell Mol Biol. 2012;47(6):738–45. Epub 20120803. doi: 10.1165/rcmb.2012-0088OC. PubMed PMID: 22865623.

22. Lau GW, Ran HM, Kong FS, Hassett DJ, Mavrodi D. Pseudomonas aeruginosa pyocyanin is critical for lung infection in mice. Infect Immun. 2004;72(7):4275–8. PubMed PMID: WOS:000222282800065.

23. Feltner JB, Wolter DJ, Pope CE, Groleau MC, Smalley NE, Greenberg EP, et al. LasR Variant Cystic Fibrosis Isolates Reveal an Adaptable Quorum-Sensing Hierarchy in Pseudomonas aeruginosa. mBio. 2016;7(5). Epub 20161004. doi: 10.1128/mBio.01513-16. PubMed PMID: 27703072; PubMed Central PMCID: PMCPMC5050340.

24. Hoffman LR, Kulasekara HD, Emerson J, Houston LS, Burns JL, Ramsey BW, Miller SI. Pseudomonas aeruginosa lasR mutants are associated with cystic fibrosis lung disease progression. J Cyst Fibros. 2009;8(1):66–70. Epub 20081029. doi: 10.1016/j.jcf.2008.09.006. PubMed PMID: 18974024; PubMed Central PMCID: PMCPMC2631641.

25. Smith EE, Buckley DG, Wu ZN, Saenphimmachak C, Hoffman LR, D’Argenio DA, et al. Genetic adaptation by Pseudomonas aeruginosa to the airways of cystic fibrosis patients. P Natl Acad Sci USA. 2006;103(22):8487–92. PubMed PMID: WOS:000238206800035.

26. Hennemann LC, LaFayette SL, Malet JK, Bortolotti P, Yang T, McKay GA, et al. LasR-deficient Pseudomonas aeruginosa variants increase airway epithelial mICAM-1 expression and enhance neutrophilic lung inflammation. PLoS Pathog. 2021;17(3):e1009375. Epub 20210310. doi: 10.1371/journal.ppat.1009375. PubMed PMID: 33690714; PubMed Central PMCID: PMCPMC7984618.

27. Aiello TB, Levy CE, Zaccariotto TR, Paschoal IA, Pereira MC, da Silva MTN, et al. Prevalence and clinical outcomes of nontuberculous mycobacteria in a Brazilian cystic fibrosis reference center. Pathog Dis. 2018;76(5). PubMed PMID: WOS:000441740900008.

28. Jonsson BE, Gilljam M, Lindblad A, Ridell M, Wold AE, Welinder-Olsson C. Molecular epidemiology of Mycobacterium abscessus, with focus on cystic fibrosis. J Clin Microbiol. 2007;45(5):1497–504. PubMed PMID: WOS:000246600300018.

29. Olivier KN, Weber DJ, Wallace RJ, Faiz AR, Lee JH, Zhang YS, et al. Nontuberculous mycobacteria I: Multicenter prevalence study in cystic fibrosis. Am J Resp Crit Care. 2003;167(6):828–34. PubMed PMID: WOS:000181439100006.

30. Sermet-Gaudelus I, Le Bourgeois M, Pierre-Audigier C, Offredo C, Guillemot D, Halley S, et al. Mycobacterium abscessus and children with cystic fibrosis. Emerg Infect Dis. 2003;9(12):1587–91. PubMed PMID: WOS:000187247600013.

31. Bernut A, Dupont C, Ogryzko NV, Neyret A, Herrmann JL, Floto RA, et al. CFTR Protects against Mycobacterium abscessus Infection by Fine-Tuning Host Oxidative Defenses. Cell Rep. 2019;26(7):1828-+. doi: 10.1016/j.celrep.2019.01.071. PubMed PMID: WOS:000458403600014.

32. Roux AL, Catherinot E, Ripoll F, Soismier N, Macheras E, Ravilly S, et al. Multicenter study of prevalence of nontuberculous mycobacteria in patients with cystic fibrosis in france. J Clin Microbiol. 2009;47(12):4124–8. Epub 20091021. doi: 10.1128/JCM.01257-09. PubMed PMID: 19846643; PubMed Central PMCID: PMCPMC2786646.

33. Byrd TF, Lyons CR. Preliminary characterization of a Mycobacterium abscessus mutant in human and murine models of infection. Infect Immun. 1999;67(9):4700–7. doi: 10.1128/IAI.67.9.4700-4707.1999. PubMed PMID: 10456919; PubMed Central PMCID: PMCPMC96797.

34. Catherinot E, Roux AL, Macheras E, Hubert D, Matmar M, Dannhoffer L, et al. Acute respiratory failure involving an R variant of Mycobacterium abscessus. J Clin Microbiol. 2009;47(1):271–4. Epub 20081119. doi: 10.1128/JCM.01478-08. PubMed PMID: 19020061; PubMed Central PMCID: PMCPMC2620830.

35. Howard ST, Rhoades E, Recht J, Pang XH, Alsup A, Kolter R, et al. Spontaneous reversion of Mycobacterium abscessus from a smooth to a rough morphotype is associated with reduced expression of glycopeptidolipid and reacquisition of an invasive phenotype. Microbiol-Sgm. 2006;152:1581–90. PubMed PMID: WOS:000238563800002.

36. Greendyke R, Byrd TF. Differential antibiotic susceptibility of Mycobacterium abscessus variants in biofilms and macrophages compared to that of planktonic bacteria. Antimicrob Agents Chemother. 2008;52(6):2019–26. Epub 20080331. doi: 10.1128/AAC.00986-07. PubMed PMID: 18378709; PubMed Central PMCID: PMCPMC2415760.

37. Roux AL, Viljoen A, Bah A, Simeone R, Bernut A, Laencina L, et al. The distinct fate of smooth and rough Mycobacterium abscessus variants inside macrophages. Open Biol. 2016;6(11). doi: 10.1098/rsob.160185. PubMed PMID: 27906132; PubMed Central PMCID: PMCPMC5133439.

38. De Boeck K, Alifier M, Vandeputte S. Sputum induction in young cystic fibrosis patients. Eur Respir J. 2000;16(1):91–4. doi: 10.1034/j.1399-3003.2000.16a16.x. PubMed PMID: 10933091.

39. Bragonzi A, Farulla I, Paroni M, Twomey KB, Pirone L, Lore NI, et al. Modelling co-infection of the cystic fibrosis lung by Pseudomonas aeruginosa and Burkholderia cenocepacia reveals influences on biofilm formation and host response. PLoS One. 2012;7(12):e52330. Epub 2013/01/04. doi: 10.1371/journal.pone.0052330. PubMed PMID: 23284990; PubMed Central PMCID: PMCPMC3528780.

40. Mangione EJ, Huitt G, Lenaway D, Beebe J, Bailey A, Figoski M, et al. Nontuberculous mycobacterial disease following hot tub exposure. Emerg Infect Dis. 2001;7(6):1039–42. doi: 10.3201/eid0706.010623. PubMed PMID: 11747738; PubMed Central PMCID: PMCPMC2631894.

41. Bjarnsholt T, Jensen PO, Fiandaca MJ, Pedersen J, Hansen CR, Andersen CB, et al. Pseudomonas aeruginosa biofilms in the respiratory tract of cystic fibrosis patients. Pediatr Pulmonol. 2009;44(6):547–58. doi: 10.1002/ppul.21011. PubMed PMID: 19418571.

42. Qvist T, Eickhardt S, Kragh KN, Andersen CB, Iversen M, Hoiby N, Bjarnsholt T. Chronic pulmonary disease with Mycobacterium abscessus complex is a biofilm infection. Eur Respir J. 2015;46(6):1823–6. PubMed PMID: WOS:000366948700040.

43. Ripoll F, Pasek S, Schenowitz C, Dossat C, Barbe V, Rottman M, et al. Non Mycobacterial Virulence Genes in the Genome of the Emerging Pathogen Mycobacterium abscessus. Plos One. 2009;4(6). doi:10.1371/journal.pone.0005660. PubMed PMID: WOS:000267237400001.

44. Adjemian J, Olivier KN, Prevots DR. Epidemiology of Pulmonary Nontuberculous Mycobacterial Sputum Positivity in Patients with Cystic Fibrosis in the United States, 2010-2014. Ann Am Thorac Soc. 2018;15(7):817–26. Epub 2018/06/14. doi: 10.1513/AnnalsATS.201709-727OC. PubMed PMID: 29897781; PubMed Central PMCID: PMCPMC6137684.

45. Hsieh MH, Lin CY, Wang CY, Fang YF, Lo YL, Lin SM, Lin HC. Impact of concomitant nontuberculous mycobacteria and Pseudomonas aeruginosa isolates in non-cystic fibrosis bronchiectasis. Infect Drug Resist. 2018;11:1137–43. Epub 20180810. doi: 10.2147/IDR.S169789. PubMed PMID: 30127630; PubMed Central PMCID: PMCPMC6089115.

46. Kamata H, Asakura T, Suzuki S, Namkoong H, Yagi K, Funatsu Y, et al. Impact of chronic Pseudomonas aeruginosa infection on health-related quality of life in Mycobacterium avium complex lung disease. Bmc Pulm Med. 2017;17(1):198. Epub 20171213. doi: 10.1186/s12890-017-0544-x. PubMed PMID: 29237500; PubMed Central PMCID: PMCPMC5727955.

47. Takano K, Shimada D, Kashiwagura S, Kamioka Y, Hariu M, Watanabe Y, Seki M. Severe Pulmonary Mycobacterium abscessus Cases Due to Co-Infection with Other Microorganisms Well Treated by Clarithromycin and Sitafloxacin in Japan. Int Med Case Rep J. 2021;14:465–70. Epub 20210712. doi: 10.2147/IMCRJ.S321969. PubMed PMID: 34285595; PubMed Central PMCID: PMCPMC8285566.

48. Idosa AW, Wozniak DJ, Hall-Stoodley L. Surface Dependent Inhibition of Mycobacterium abscessus by Diverse Pseudomonas aeruginosa Strains. Microbiol Spectr. 2022:e0247122. Epub 20221117. doi: 10.1128/spectrum.02471-22. PubMed PMID: 36394312.

49. Rodriguez-Sevilla G, Crabbe A, Garcia-Coca M, Aguilera-Correa JJ, Esteban J, Perez-Jorge C. Antimicrobial Treatment Provides a Competitive Advantage to Mycobacterium abscessus in a Dual-Species Biofilm with Pseudomonas aeruginosa. Antimicrob Agents Chemother. 2019;63(11). Epub 20191022. doi: 10.1128/AAC.01547-19. PubMed PMID: 31451500; PubMed Central PMCID: PMCPMC6811427.

50. Rodriguez-Sevilla G, Garcia-Coca M, Romera-Garcia D, Aguilera-Correa JJ, Mahillo-Fernandez I, Esteban J, Perez-Jorge C. Non-Tuberculous Mycobacteria multispecies biofilms in cystic fibrosis: development of an in vitro Mycobacterium abscessus and Pseudomonas aeruginosa dual species biofilm model. Int J Med Microbiol. 2018;308(3):413–23. PubMed PMID: WOS:000443629800010.

51. Bajorath J, Hinrichs W, Saenger W. The enzymatic activity of proteinase K is controlled by calcium. Eur J Biochem. 1988;176(2):441–7. doi: 10.1111/j.1432-1033.1988.tb14301.x. PubMed PMID: 3166426.

52. Vrla GD, Esposito M, Zhang C, Kang Y, Seyedsayamdost MR, Gitai Z. Cytotoxic alkyl-quinolones mediate surface-induced virulence in Pseudomonas aeruginosa. PLoS Pathog. 2020;16(9):e1008867. Epub 20200914. doi: 10.1371/journal.ppat.1008867. PubMed PMID: 32925969; PubMed Central PMCID: PMCPMC7515202.

53. Toung JM, Morley M, Li MY, Cheung VG. RNA-sequence analysis of human B-cells. Genome Res. 2011;21(6):991–8. doi: 10.1101/gr.116335.110. PubMed PMID: WOS:000291153400019.

54. Westermann AJ, Forstner KU, Amman F, Barquist L, Chao YJ, Schulte LN, et al. Dual RNA-seq unveils noncoding RNA functions in host-pathogen interactions. Nature. 2016;529(7587):496-+. doi: 10.1038/nature16547. PubMed PMID: WOS:000368673800030.

55. Beste DJV, Espasa M, Bonde B, Kierzek AM, Stewart GR, McFadden J. The Genetic Requirements for Fast and Slow Growth in Mycobacteria. Plos One. 2009;4(4). doi: 10.1371/journal.pone.0005349. PubMed PMID: WOS:000265514900012.

56. Maurya RK, Bharti S, Krishnan MY. Triacylglycerols: Fuelling the Hibernating Mycobacterium tuberculosis. Front Cell Infect Microbiol. 2018;8:450. Epub 20190109. doi: 10.3389/fcimb.2018.00450. PubMed PMID: 30687647; PubMed Central PMCID: PMCPMC6333902.

57. Shi L, Sohaskey CD, Kana BD, Dawes S, North RJ, Mizrahi V, Gennaro ML. Changes in energy metabolism of Mycobacterium tuberculosis in mouse lung and under in vitro conditions affecting aerobic respiration. Proc Natl Acad Sci U S A. 2005;102(43):15629–34. Epub 20051014. doi: 10.1073/pnas.0507850102. PubMed PMID: 16227431; PubMed Central PMCID: PMCPMC1255738.

58. Dedrick RM, Aull HG, Jacobs-Sera D, Garlena RA, Russell DA, Smith BE, et al. The Prophage and Plasmid Mobilome as a Likely Driver of Mycobacterium abscessus Diversity. Mbio. 2021;12(2). doi: 10.1128/mBio.03441-20. PubMed PMID: WOS:000643684900005.

59. Miao Zhao KG, Lia Danelishvili , Brendan Jeffrey, Luiz E Bermudez. Identification of Prophages within the Mycobacterium avium 104 Genome and the Link of Their Function Regarding to Environment Survival. Adv Microbiol. 2016;6(13):927–41. PubMed Central PMCID: PMCPMC8293804.

60. van Dijk D, Dhar R, Missarova AM, Espinar L, Blevins WR, Lehner B, Carey LB. Slow-growing cells within isogenic populations have increased RNA polymerase error rates and DNA damage. Nat Commun. 2015;6. doi: 10.1038/ncomms8972. PubMed PMID: WOS:000360346900013.

61. Munoz-Elias EJ, Upton AM, Cherian J, McKinney JD. Role of the methylcitrate cycle in Mycobacterium tuberculosis metabolism, intracellular growth, and virulence. Mol Microbiol. 2006;60(5):1109–22. doi: 10.1111/j.1365-2958.2006.05155.x. PubMed PMID: 16689789.

62. Savvi S, Warner DF, Kana BD, McKinney JD, Mizrahi V, Dawes SS. Functional characterization of a vitamin B12-dependent methylmalonyl pathway in Mycobacterium tuberculosis: implications for propionate metabolism during growth on fatty acids. J Bacteriol. 2008;190(11):3886–95. Epub 20080328. doi: 10.1128/JB.01767-07. PubMed PMID: 18375549; PubMed Central PMCID: PMCPMC2395058.

63. Viljoen A, Blaise M, de Chastellier C, Kremer L. MAB_3551c encodes the primary triacylglycerol synthase involved in lipid accumulation in Mycobacterium abscessus. Mol Microbiol. 2016;102(4):611–27. doi: 10.1111/mmi.13482. PubMed PMID: WOS:000387757100005.

64. Sezonov G, Joseleau-Petit D, D’Ari R. Escherichia coli physiology in Luria-Bertani broth. J Bacteriol. 2007;189(23):8746–9. Epub 20070928. doi: 10.1128/JB.01368-07. PubMed PMID: 17905994; PubMed Central PMCID: PMCPMC2168924.

65. Garton NJ, Waddell SJ, Sherratt AL, Lee SM, Smith RJ, Senner C, et al. Cytological and transcript analyses reveal fat and lazy persister-like bacilli in tuberculous sputum. PLoS Med. 2008;5(4):e75. doi: 10.1371/journal.pmed.0050075. PubMed PMID: 18384229; PubMed Central PMCID: PMCPMC2276522.

66. Russell DG, Cardona PJ, Kim MJ, Allain S, Altare F. Foamy macrophages and the progression of the human tuberculosis granuloma. Nat Immunol. 2009;10(9):943–8. Epub 20090819. doi: 10.1038/ni.1781. PubMed PMID: 19692995; PubMed Central PMCID: PMCPMC2759071.

67. Biswas L, Gotz F. Molecular Mechanisms of Staphylococcus and Pseudomonas Interactions in Cystic Fibrosis. Front Cell Infect Microbiol. 2021;11:824042. Epub 20220106. doi: 10.3389/fcimb.2021.824042. PubMed PMID: 35071057; PubMed Central PMCID: PMCPMC8770549.

68. Lamont IL, Konings AF, Reid DW. Iron acquisition by Pseudomonas aeruginosa in the lungs of patients with cystic fibrosis. Biometals. 2009;22(1):53–60. Epub 20090107. doi: 10.1007/s10534-008-9197-9. PubMed PMID: 19130260.

69. Martin LW, Reid DW, Sharples KJ, Lamont IL. Pseudomonas siderophores in the sputum of patients with cystic fibrosis. Biometals. 2011;24(6):1059–67. Epub 20110604. doi: 10.1007/s10534-011-9464-z. PubMed PMID: 21643731.

70. Lin J, Cheng J, Wang Y, Shen X. The Pseudomonas Quinolone Signal (PQS): Not Just for Quorum Sensing Anymore. Front Cell Infect Microbiol. 2018;8:230. Epub 20180704. doi: 10.3389/fcimb.2018.00230. PubMed PMID: 30023354; PubMed Central PMCID: PMCPMC6039570.

71. Stintzi A, Evans K, Meyer JM, Poole K. Quorum-sensing and siderophore biosynthesis in Pseudomonas aeruginosa: lasR/lasI mutants exhibit reduced pyoverdine biosynthesis. FEMS Microbiol Lett. 1998;166(2):341–5. doi: 10.1111/j.1574-6968.1998.tb13910.x. PubMed PMID: 9770291.

72. Venturi V. Regulation of quorum sensing in Pseudomonas. FEMS Microbiol Rev. 2006;30(2):274–91. doi: 10.1111/j.1574-6976.2005.00012.x. PubMed PMID: 16472307.

73. Birmes FS, Saring R, Hauke MC, Ritzmann NH, Drees SL, Daniel J, et al. Interference with Pseudomonas aeruginosa Quorum Sensing and Virulence by the Mycobacterial Pseudomonas Quinolone Signal Dioxygenase AqdC in Combination with the N-Acylhomoserine Lactone Lactonase QsdA. Infect Immun. 2019;87(10). Epub 20190919. doi: 10.1128/IAI.00278-19. PubMed PMID: 31308081; PubMed Central PMCID: PMCPMC6759295.

74. Letoffe S, Wu Y, Darch SE, Beloin C, Whiteley M, Touqui L, Ghigo JM. Pseudomonas aeruginosa Production of Hydrogen Cyanide Leads to Airborne Control of Staphylococcus aureus Growth in Biofilm and In Vivo Lung Environments. mBio. 2022;13(5):e0215422. Epub 2022/09/22. doi: 10.1128/mbio.02154-22. PubMed PMID: 36129311; PubMed Central PMCID: PMCPMC9600780.

75. Nolan LM, Allsopp LP. Antimicrobial Weapons of Pseudomonas aeruginosa. Adv Exp Med Biol. 2022;1386:223–56. Epub 2022/10/19. doi: 10.1007/978-3-031-08491-1_8. PubMed PMID: 36258074.

76. Holloway BW, Krishnapillai V, Morgan AF. Chromosomal genetics of Pseudomonas. Microbiol Rev. 1979;43(1):73–102. doi: 10.1128/mr.43.1.73-102.1979. PubMed PMID: 111024; PubMed Central PMCID: PMCPMC281463.

77. Wilder CN, Diggle SP, Schuster M. Cooperation and cheating in Pseudomonas aeruginosa: the roles of the las, rhl and pqs quorum-sensing systems. ISME J. 2011;5(8):1332–43. Epub 20110303. doi: 10.1038/ismej.2011.13. PubMed PMID: 21368905; PubMed Central PMCID: PMCPMC3146268.

78. Howard ST, Byrd TF. The rapidly growing mycobacteria: saprophytes and parasites. Microbes Infect. 2000;2(15):1845–53. PubMed PMID: WOS:000166719500008.

79. Gupta R, Netherton M, Byrd TF, Rohde KH. Reporter-Based Assays for High-Throughput Drug Screening against Mycobacterium abscessus. Front Microbiol. 2017;8:2204. Epub 20171110. doi: 10.3389/fmicb.2017.02204. PubMed PMID: 29176967; PubMed Central PMCID: PMCPMC5687050.

80. Costa KC, Bergkessel M, Saunders S, Korlach J, Newman DK. Enzymatic Degradation of Phenazines Can Generate Energy and Protect Sensitive Organisms from Toxicity. Mbio. 2015;6(6). PubMed PMID: WOS:000367524700024.

81. Newman JR, Fuqua C. Broad-host-range expression vectors that carry the L-arabinose-inducible Escherichia coli araBAD promoter and the araC regulator. Gene. 1999;227(2):197–203. doi: 10.1016/s0378-1119(98)00601-5. PubMed PMID: 10023058.

82. Lee JH, Lequette Y, Greenberg EP. Activity of purified QscR, a Pseudomonas aeruginosa orphan quorum-sensing transcription factor. Mol Microbiol. 2006;59(2):602–9. doi: 10.1111/j.1365-2958.2005.04960.x. PubMed PMID: 16390453.

83. Wilder CN, Allada G, Schuster M. Instantaneous within-patient diversity of Pseudomonas aeruginosa quorum-sensing populations from cystic fibrosis lung infections. Infect Immun. 2009;77(12):5631–9. Epub 20091005. doi: 10.1128/IAI.00755-09. PubMed PMID: 19805523; PubMed Central PMCID: PMCPMC2786440.

84. Rohde KH, Veiga DFT, Caldwell S, Balazsi G, Russell DG. Linking the Transcriptional Profiles and the Physiological States of Mycobacterium tuberculosis during an Extended Intracellular Infection. Plos Pathogens. 2012;8(6). PubMed PMID: WOS:000305987800039.

85. Kim D, Paggi JM, Park C, Bennett C, Salzberg SL. Graph-based genome alignment and genotyping with HISAT2 and HISAT-genotype. Nat Biotechnol. 2019;37(8):907-+. PubMed PMID: WOS:000482876100022.

86. Liao Y, Smyth GK, Shi W. featureCounts: an efficient general purpose program for assigning sequence reads to genomic features. Bioinformatics. 2014;30(7):923–30. PubMed PMID: WOS:000334078300005.

87. Ritchie ME, Phipson B, Wu D, Hu Y, Law CW, Shi W, Smyth GK. limma powers differential expression analyses for RNA-sequencing and microarray studies. Nucleic Acids Res. 2015;43(7):e47. Epub 20150120. doi: 10.1093/nar/gkv007. PubMed PMID: 25605792; PubMed Central PMCID: PMCPMC4402510.

